# H3K9me3 is Required for Inheritance of Small RNAs that Target a Unique Subset of Newly Evolved Genes

**DOI:** 10.1101/338582

**Authors:** Itamar Lev, Hila Gingold, Oded Rechavi

## Abstract

In *Caenorhabditis elegans*, RNA interference (RNAi) responses can transmit across generations via small RNAs. RNAi inheritance is associated with Histone-3-Lysine-9 tri-methylation (H3K9me3) of the targeted genes. In other organisms, maintenance of silencing requires a feed-forward loop between H3K9me3 and small RNAs. Here we show that in *C. elegans* not only is H3K9me3 unnecessary for inheritance, the modification’s function depends on the identity of the RNAi-targeted gene. We found an asymmetry in the requirement for H3K9me3 and the main worm H3K9me3 methyltransferases, SET-25 and SET-32. Both methyltransferases promote heritable silencing of the foreign gene *gfp*, but are dispensable for silencing of the endogenous gene *oma-1*. Genome-wide examination of heritable endogenous small interfering RNAs (endo-siRNAs) revealed that the SET-25-dependent heritable endo-siRNAs target newly acquired and highly H3K9me3 marked genes. Thus, “repressive” chromatin marks could be important specifically for heritable silencing of genes which are flagged as “foreign”, such as *gfp*.

## Introduction

RNA interference (RNAi) responses are inherited in *Caenorhabditis elegans* nematodes across generations via heritable small RNAs (Rechavi & Lev, 2017). In worms, exposure to a number of environmental challenges, such as viral infection (Rechavi *et al*, 2011; Gammon *et al*, 2017), starvation (Rechavi *et al*, 2014), and heat (Klosin *et al*, 2017) induces heritable physiological responses that persist for multiple generations. Inheritance of such transmitted information was linked to inheritance of small RNAs and chromatin modifications, and hypothesized to protect and prepare the progeny for the environmental challenges that the ancestors met.

By base-pairing with complementary mRNA sequences, small RNAs in *C. elegans* control the expression of thousands of genes, and protect the genome from foreign elements (Malone & Hannon, 2009; Luteijn & Ketting, 2013). Via recruitment of RNA-binding proteins, small interfering RNAs (siRNAs) can induce gene silencing also by inhibiting transcription (Castel & Martienssen, 2013).

Small RNA-mediated transcription inhibition involves modification of histones, however the exact role that histone marks play in inheritance of RNAi and small RNA synthesis is under debate (Rechavi & Lev, 2017). Worms small RNAs that enter the nucleus were shown to inhibit the elongation phase of Pol II (Guang *et al*, 2010); In addition, nuclear small RNAs are thought to recruit histone modifiers to the target’s chromatin, resulting in deposition of histone marks such as histone H3K9-tri methylation (H3K9me3) and H3K27me3 (Gu *et al*, 2012; Mao *et al*, 2015; Lev *et al*, 2017).

The interaction between small RNAs and repressive chromatin marks are reciprocal: in *Arabidopsis thaliana* (Molnar *et al*, 2010; Daniel Holoch *et al*, 2015) and *Schizosaccharomyces pombe* (Verdel *et al*, 2004; Moazed *et al*, 2006), small RNAs and repressive histone marks form a self-reinforcing feed-forward loop, where nuclear small RNAs induce deposition of repressive histone marks, and in turn the repressive chromatin marks recruit the small RNA machinery to synthesize additional small RNAs. Whether a similar feedback operates in worms and other organisms, is unclear. In *Neurospora crassa*, transgene-induced small RNAs work independently of H3K9me3 (Chicas *et al*, 2005). In *C. elegans*, it was previously suggested that H3K9me is required for RNAi inheritance (Shirayama *et al*, 2012). However, studies from different groups have shown that the situation is more complex, and that H3K9me could be dispensable, and can even suppress heritable silencing of some targets (Kalinava *et al*, 2017; Lev *et al*, 2017; Minkina & Hunter, 2017).

In *C. elegans* H3K9me is considered to depend mainly on the methyltransferases MET-2, SET-25, and SET-32 (Towbin *et al*, 2012; Kalinava *et al*, 2017; Spracklin *et al*, 2017). H3K9 methylation by MET-2 and SET-25 occurs in a step-wise fashion – after MET-2 deposits the first two methyl groups (H3K9me1/2), SET-25 can add the third methyl group (me3) (Towbin *et al*, 2012). In the germline, however, SET-25 is capable of tri-methylating H3K9 in a MET-2-independent manner (Bessler *et al*, 2010; Towbin *et al*, 2012). SET-32-dependent H3K9me3 is at least in part independent of the activity of SET-25 or MET-2 (Kalinava *et al*, 2017).

To study the roles of H3K9me3 in the maintenance of heritable small RNAs we examined the inheritance of small RNAs in mutants defective in these histone methyltransferases. Although H3K9me3 was thought to be required for heritable RNAi (Ashe *et al*, 2012; Gu *et al*, 2012), we found the heritable RNAi-responses are greatly potentiated in *met-2* mutant background (Lev *et al*, 2017). Our data indicated that the enhanced strength of the RNAi responses in *met-2* mutants stems from a genome-wide massive loss of different endogenous small RNA (endo-siRNAs) species. In normal circumstances, these endo-siRNAs compete with exogenously derived siRNAs over shared biosynthesis components required for small RNA production or inheritance (Lev *et al*, 2017). In addition, we found that the accumulated sterility (or “Mortal Germline”, Mrt phenotype) of *met-2* mutants results from dysfunctional small RNA inheritance (Lev *et al*, 2017).

However, our previous results regarding the role of H3K9me1/2 (deposited by MET-2) did not rule out the possibility that H3K9me3 is yet required for efficient heritable silencing of *gfp* transgenes: We found that RNAi responses in *met-2* mutants nevertheless lead to marking of the target gene’s histones with a heritable H3K9me3 modification. Further, a comparison of the H3K9me3 signal on the *gfp* locus in different mutants has shown that anti-*gfp* RNAi responses were strongly inherited only in genetic backgrounds where some H3K9me3 trace could be detected (i.e, in wild type, *met-2,* and *met-2;set-25* mutants). In *set-25* single mutants, where no statistically significant H3K9me3 footprint could be detected, anti-*gfp* RNAi was only weakly inherited. Previously, *set-25* mutants were reported to be deficient in heritable RNAi responses targeting different fluorescent transgenes (Ashe *et al*, 2012; Lev *et al*, 2017; Spracklin *et al*, 2017).

In contrast to anti-*gfp* heritable RNAi responses, for which H3K9me3 is important, we detected an enhancement in the inheritance potency of anti-*oma-1* RNAi in *set-25* mutants (Lev *et al*, 2017). However, in that study we did not examine whether an H3K9me3 footprint was deposited on the endogenous gene *oma-1* in the *set-25* background (Lev *et al*, 2017). The publication of a recent paper (Kalinava *et al*, 2017) which described strong anti-*oma-1* RNAi inheritance in *met-2;set-25;set-32* triple mutants, despite the absence of a detectable H3K9me3 footprint, prompted us to re-examine the inheritance of anti-*gfp* RNAi in this triple mutants. We hypothesized that gene-specific characteristics lead to contrasting requirements for H3K9me3 and specific methyltransferases. In this manuscript we describe an asymmetry in the requirement for H3K9me3 and specific methyltransferases for heritable RNAi responses aimed against the endogenous gene *oma-1* and the foreign gene *gfp*. These differences led us to perform a genome-wide analysis of SET-25-dependent small RNAs, which revealed that the endo-siRNAs which depend on H3K9me3 target newly acquired *C. elegans* genes that might be considered “foreign”, similarly to *gfp*.

## Results

Recently Kalinava et al. examined the heritable RNAi responses against *oma-1* also in a triple mutant, lacking the three main *C. elegans* H3K9 methyltransferases, SET-25, SET-32 and MET-2 (Kalinava *et al*, 2017). The authors reported that silencing of *oma-1* was independent of H3K9me3, as in these mutants RNAi responses raised against the *oma-1* gene were heritable despite the lack of an H3K9me3 trace (Kalinava *et al*, 2017).

We successfully replicated the results of Kalinava et al., and came to the same conclusion, that the *met-2;set-25;set-32* triple mutant worms inherit RNAi responses against the *oma-1* gene, also when we used a different assay for inheritance (**Figure 1A** and **Figure 1B, upper panel**). Unlike Kalinava et al., which used qPCR to score for downregulation of *oma-1* expression, we targeted a redundant, temperature-sensitive *oma-1* allele, that in the restrictive temperatures does not allow the development of embryos unless silenced (as previously described (Alcazar *et al*, 2008)). Upon shifting to 20 degrees, only worms that silence the *oma-1* gene in a heritable manner survive.

**Figure 1.**
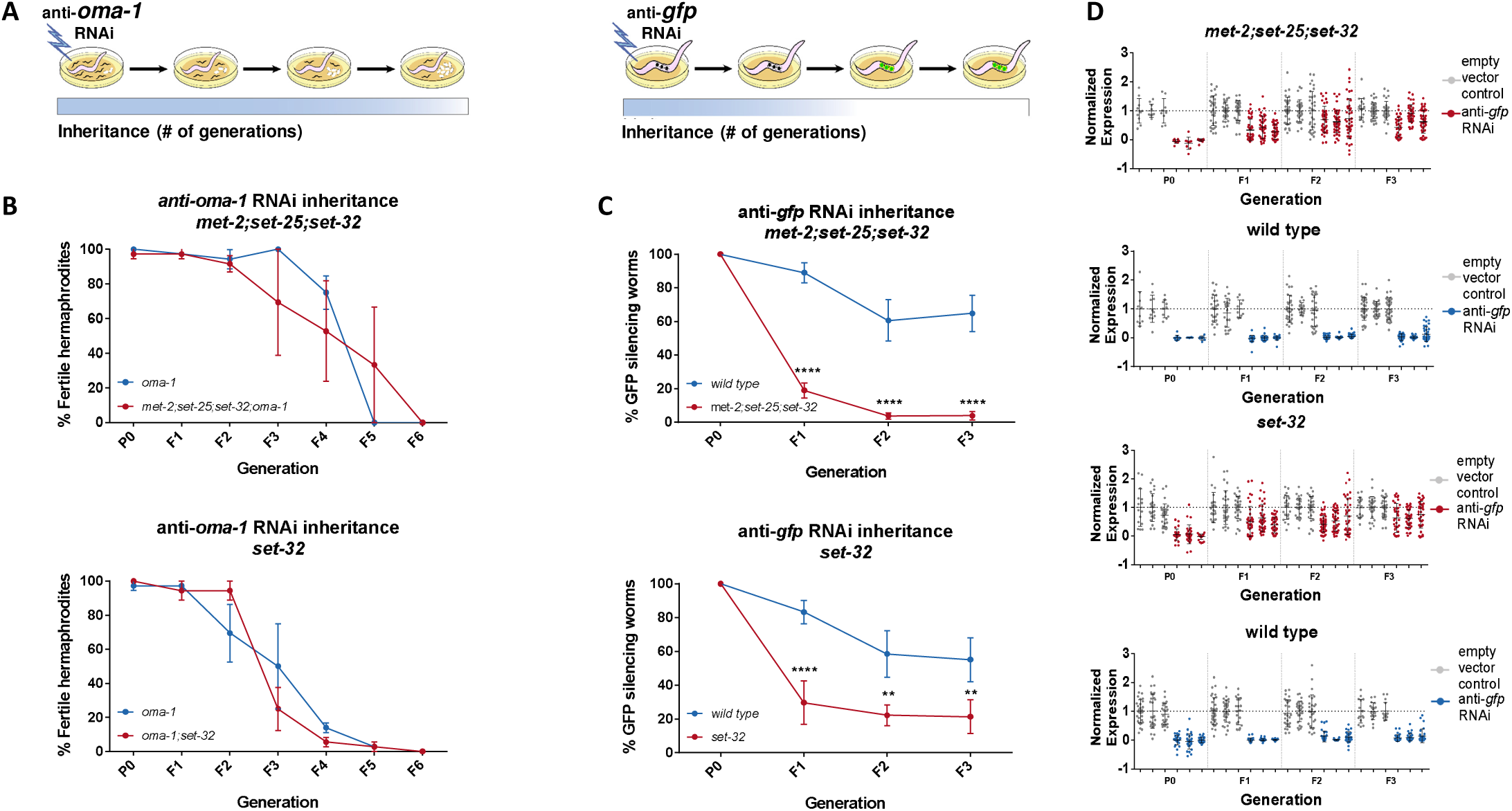
Heritable RNAi responses against the *oma-1* and *gfp* genes have different requirements for H3K9me3 methyltransferases. **(A)** Scheme depicting the different requirements for H3K9 methyltransferases in RNAi inheritance responses aimed at different genes. (left) Only worms that inherit small RNAs that silence the temperature-sensitive dominant allele of *oma-1* can hatch. Heritable RNAi responses aimed against the endogenous *oma-1* gene do not require H3K9me3 methyltransferases. (right) Inheritance of anti-*gfp* small RNAs lead to heritable silencing of the *gfp* transgene. Heritable RNAi responses aimed against the foreign *gfp* gene strongly depends on H3K9me3 methyltransferases. **(B)** Inheritance of anti-*oma-*1 RNAi response in H3K9me3 methyltransferase mutants. The percentage of fertile worms per replicate and generation is presented (N = 12, three biological replicates). (upper panel) RNAi inheritance dynamics in *met-2;set-25;set-32;oma-1* mutants compared to *oma-1* mutants. (lower panel) RNAi inheritance dynamics in *set-32* single mutants compared to wild type. **(C)** Inheritance of anti-*gfp* RNAi response in H3K9me3 methyltransferase mutants. In each generation the percentage of worms silencing a germline expressed GFP transgene is presented (N >60, three replicates). (upper panel) RNAi inheritance dynamics in *met-2;set-25;set-32* triple mutants. (lower panel) RNAi inheritance dynamics in *set-32* single mutants. Error bars represent standard error of mean. *p-value < 0.05, **p-value < 0.005, ***p-value < 0.001, ****p-value < 0.0001, Two-way ANOVA, Sidak’s multiple comparisons test. **(D)** Quantification of the germline GFP expression levels in RNAi treated animals normalized to untreated animals. (upper panel) GFP silencing Inheritance dynamics in *met-2;set-25;set-32* triple mutants (Red) and wild type animals (Blue, In total, the germlines of **1350** worms were quantified). (lower panel) GFP silencing Inheritance dynamics in *set-32* mutants (Red) and wild type animals (Blue, in total, the germlines of **1480** worms were quantified). Error bars represent standard deviation.

In parallel we discovered, surprisingly, that in contrast to anti-*oma-1* inheritance, heritable silencing of a *gfp* transgene was defective in the same triple mutants (**Figure 1C, upper panel**, p=0.0014, 2-way ANOVA). In addition, we also confirmed (Spracklin *et al*, 2017) that while *set-32* single mutants are deficient in inheriting RNAi responses raised against the *gfp* transgene (**Figure 1C, lower panel,** p=0.0026, 2-way ANOVA), they are capable (Kalinava *et al*, 2017) of inheriting responses raised against *oma-1* (**Figure 1B, lower panel,** p=0.8487, 2-way ANOVA). Previously we have shown that while *set-25* mutants are defective in inheritance of anti-*gfp* RNAi, weak inheritance responses can still be observed (Lev *et al*, 2017). Similarly, we were able to detect weak inheritance responses that last at least until the F3 generation also in *met-2;set-25;set-32* and *set-32* mutants (**Figure 1D**, p-value < 0.0001 for *met-2;set-25;set-32* and *set-32* in the F3 generation, Two-way ANOVA). Together with our previous data, which showed that *set-25* is required for inheriting anti-*gfp* RNAi, but not anti-*oma-1* RNAi (Lev *et al*, 2017), these results suggested that heritable RNAi requires H3K9 methyltransferases in a gene-specific manner.

### The levels of RNAi-induced H3K9me3 do not explain the gene-specific requirements of methyltransferases for heritable RNAi

Histone methyltransferase mutants may affect RNAi-induced H3K9me3 levels in a gene-specific manner, thus leading to different inheritance dynamics for each gene. To test this possibility, we performed anti-H3K9me3 Chromatin Immunoprecipitation (ChIP) on F1 *met-2;set-25;set-32* triple mutant progeny, that were derived from parents exposed to anti-*oma-1* RNAi, anti-*gfp* RNAi, or untreated controls. Using qPCR we found, as expected (Kalinava *et al*, 2017) that in *met-2;set-25;set-32* triple mutants the RNAi-induced H3K9me3 signal was significantly reduced (p-value = 0.0007 and 0.0009, Two-way ANOVA, for *gfp* and *oma-1*, respectively). Importantly, this was true for both the *oma-1* and *gfp* loci (**Figure 2A**). Interestingly, in naive wild-type animals, that were not treated with RNAi, the levels of H3K9me3 on *gfp* were significantly higher than on *oma-1* (**Figure 2B**, p-value = 0.0039). Regardless, as no differences can be found in the RNAi-induced fold changes in H3K9me3 levels between *gfp* and *oma-1* (**Figure 2A**), the levels of *RNAi-induced* H3K9me3 cannot explain the gene-specific requirements of methyltransferases for heritable RNAi.

**Figure 2.**
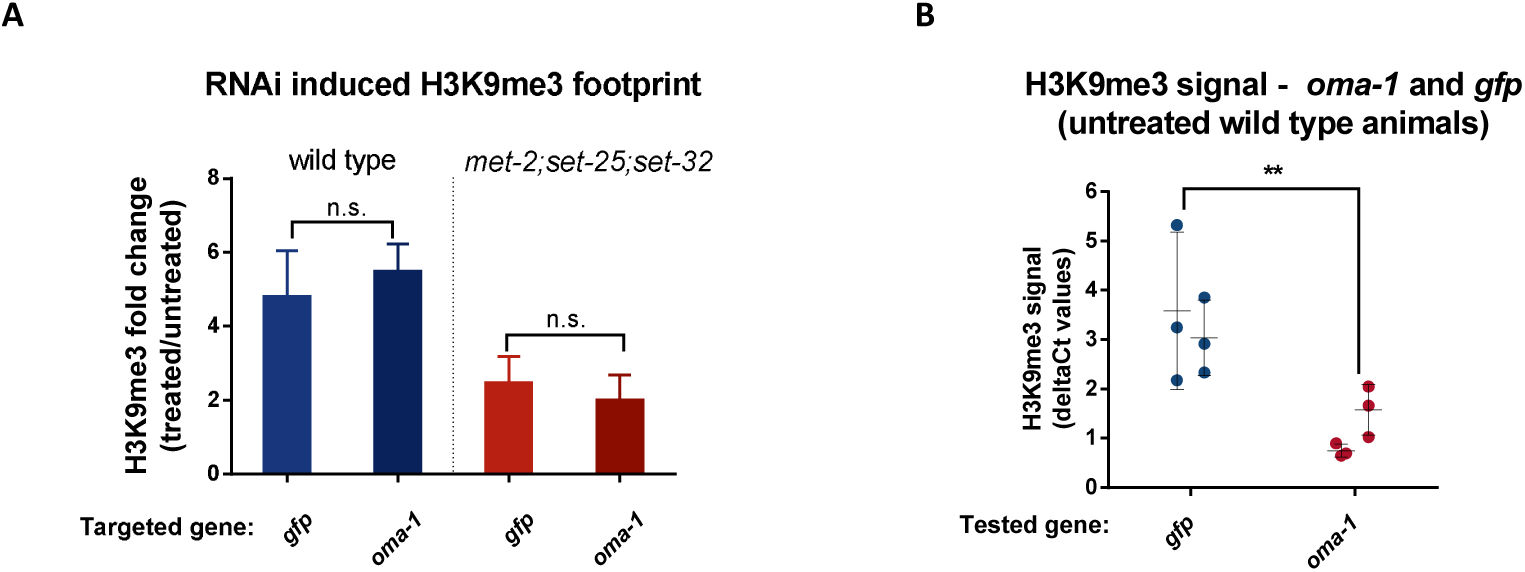
The change in RNAi-induced H3K9me3 on *oma-1* and *gfp* is comparable. **(A)** The RNAi-induced H3K9me3 footprint on the RNAi-targeted genes. The fold change in H3K9me3 levels in F1 progeny of animals exposed to RNAi versus untreated control animals. The H3K9me3 footprint levels were assessed using a qPCR quantification of ChIP experiments conducted with both wild type (left) and *met-2;set-25;set-32* mutants (right). **(B)** H3K9me3 levels on the *gfp* and *oma-1* genes in naive untreated wild type animals. The H3K9me3 signal on each of the investigated loci was measured using two different primer sets presented as filled circles (primer set #1) and empty circles (primer set #2). The deltaCt numbers used to obtain the fold change values were calculated using the *eft-3* gene as an endogenous control. The presented data was obtained from three biological replicates. The levels of *gfp* H3K9me3 signal in wild type animals are adapted from our previous publication (Lev *et al*, 2017). Two-way ANOVA, Sidak’s multiple comparisons test. *p-value<0.05, **p-value<0.005, ***p-value<0.001, ****p-value<0.0001. Error bars represent standard deviations.

### SET-32 acts upstream to MET-2 and SET-25 to support RNAi inheritance

We previously found that in contrast to *set-25* single mutants, which are deficient in RNAi-induced heritable H3K9me3 methylation (Mao *et al*, 2015; Lev *et al*, 2017), *met-2;set-25* double mutants display a modest but robust H3K9me3 footprint following RNAi (Lev *et al*, 2017; Kalinava *et al*, 2017). We therefore hypothesized that in the *met-2* background, an additional, perhaps otherwise inactive H3K9 methyltransferase, is expressed or activated, compensating for the absence of SET-25, to allow efficient heritable RNAi responses. To test this hypothesis, we first examined whether *met-2;set-32* double mutants can inherit RNAi responses raised against *gfp.* If SET-32 and SET-25 compensate for each other and are redundant, then *met-2;set-32* double mutants are expected to strongly inherit RNAi responses, similar to *met-2;set-25* double mutants (Lev *et al*, 2017). Our results show, that in contrast to *met-2;set-25* double mutants, *met-2;set-32* double mutants are defective in RNAi inheritance raised against *gfp*, since only a very weak response can be detected (**Figure 3**). The potency of RNAi inheritance in *met-2;set-32* double mutants is comparable to that of *set-25* (Lev *et al*, 2017) and *set-32* single mutants, or *met-25;set-25;set-32* triple mutants (**Figure 1C**). These results suggest that SET-32 has a distinct role, and that it probably acts upstream to MET-2 and SET-25, in promoting RNAi inheritance. This conclusion is also consistent with the recent observation that SET-32, in contrast to MET-2 and SET-25 has an essential role in establishment of RNAi-mediated nuclear silencing (preprint: (Kalinava *et al*, 2018)).

**Figure 3.**
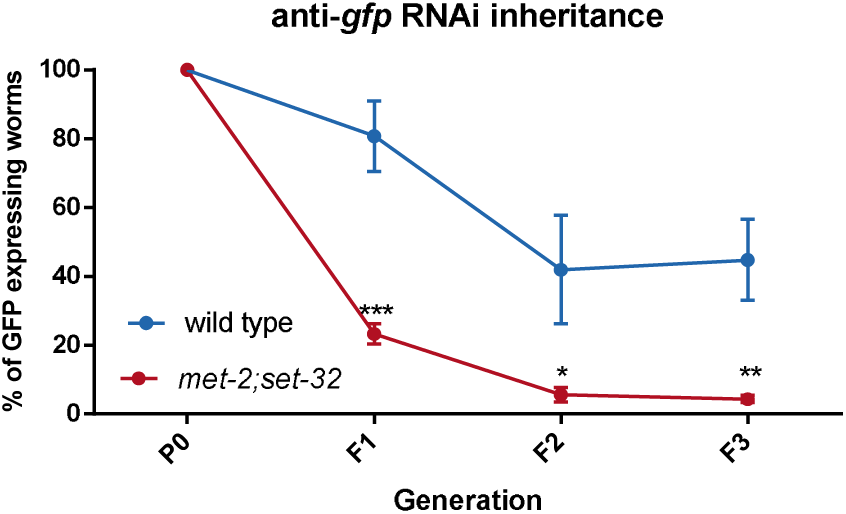
SET-32 is required for strong heritable RNAi-induced silencing of *gfp* in *met-2* mutants. Inheritance of anti-*gfp* RNAi response in *set-32;met-2* double mutants and wild type is shown. In each generation, the percentage of worms silencing a germline expressed GFP transgene is presented (N > 60, three biological replicates). *p-value<0.05, **p-value<0.005, ***p-value<0.001, Two-way ANOVA, Sidak’s multiple comparisons test. Error bars represent standard error of mean.

### SET-25 is required for the maintenance of a specific class of endo-siRNAs

Certain germline small RNAs have evolved to confer immunity against foreign genetic elements, while sparing endogenous genes (Malone & Hannon, 2009). The different requirements for particular methyltransferase and H3K9me3 for heritable silencing of *gfp* and *oma-1* may be connected to the fact that *gfp* is a “foreign” gene, while *oma-1* is an endogenous gene. We found that exogenous siRNAs that target *gfp* are lost in *set-25* mutants, and hypothesized that endo-siRNAs that target other “foreign” genes would be likewise affected. Therefore, we re-analyzed our previously published small RNA sequencing data, obtained from *set-25* mutants (Lev *et al*, 2017). However, among the targets of these differentially expressed endo-siRNAs, we could not detect striking changes (fold change >1.2) in endo-siRNAs that target transposons and repetitive elements in *set-25* mutants (**Figure 4A, left panel**). In contrast, a subset of endo-siRNAs that target 279 different protein-coding genes was found to exhibit significant changes in *set-25* mutants (**Figure 4A, right panel**). To understand why these small RNAs are uniquely affected by SET-25, we characterized this group of target genes and the endo-siRNAs that target them.

**Figure 4.**
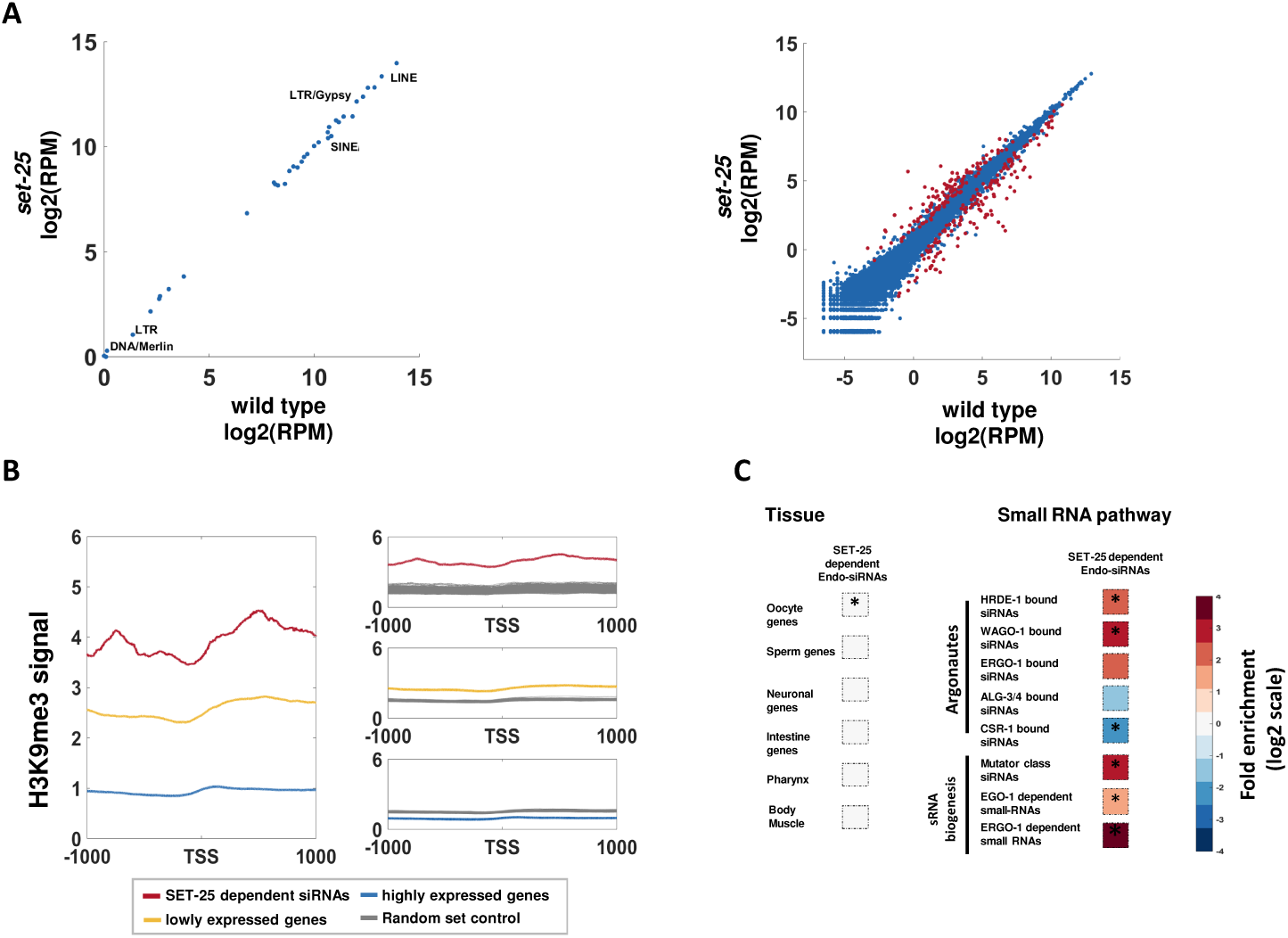
Genome-wide analyses of SET-25-dependent endo-siRNAs. (**A**) An expression analysis of endo-siRNAs targeting transposons and repetitive elements classes (left panel) or protein-coding genes (right panel). Shown are the expression values (log2 of RPM) in *set-25* mutants (y-axis) compared to wild type worms (x-axis). endo-siRNAs which display significant differential expression (analyzed with Deseq2, adjusted p-value < 0.1) are marked in red An analysis of H3K9me3 signals (based on published data from (McMurchy *et al*, 2017)) on highly expressed genes (top 10%, Blue), lowly expressed genes (top 10%, Yellow) and gene targets of SET-25 dependent endo-siRNAs (based on (Lev *et al*, 2017), Red). All genes are aligned according to their Transcription Start Sites (TSS), and the regions of 1000 base pairs upstream and downstream of the TSS are shown on the x axis. The y axis shows the averaged signal of the H3K9me3 modification as a function of distance from the TSS. An enrichment analysis of genes with significantly lowered levels of endo-siRNAs targeting them in *set-25* mutants compared to wild type. The enrichment values for expression in specific tissues (left panel) and the enrichment values for different small RNA pathways (right panel) are presented. Fold enrichment values are color coded. The p-values were calculated using 10,000 random gene sets identical in their size to the set of SET-25-dependent endo-siRNAs. Asterisk denotes statistically significant enrichment values (p-value < 0.05).

Since in *set-25* the loss of exogenous siRNAs coincided with the loss of heritable RNAi-induced H3K9me3 (Lev *et al*, 2017), we first tested whether genes that were differentially targeted by endo-siRNAs in *set-*25 mutants were also marked by H3K9me3. By examining publicly available H3K9me3 data (McMurchy *et al*, 2017), we found that the 151 genes that lost the endo-siRNAs that target them in *set-25* mutants were robustly marked by H3K9me3 in wild type animals (**Figure 4B)**. In contrast, the 128 genes that had *increased* endo-siRNA levels that target them in *set-25* mutants were not significantly marked by H3K9me3 (**Figure S1A**). These results support the hypothesis that when it comes to the 151 genes targeted by SET-25 dependent endo-siRNAs, SET-25 affects endo-siRNA biogenesis by tri-methylating H3K9.

Next we examined whether genes that display altered endo-siRNAs levels in *set-25* mutants are expressed in specific tissues. Genes that had significantly reduced levels of endo-siRNAs targeting them in *set-25* mutants, exhibited significant, but modest, enrichment for expression in the germline, specifically in oocytes (fold enrichment = 1.24, p-value = 0.0111, **Figure 4C** and **Figure S1B**). No significant enrichment was found for other tissues (**Figure 4C)**.

To identify the small RNA pathways which are affected by *set-25*, we tested whether the differentially expressed endo-siRNAs depend on particular argonautes, or associate with specific biosynthesis or functional pathways (**Figure 4C**). It was previously suggested that the CSR-1 argonaute carries heritable endo-siRNAs that mark endogenous genes (Claycomb *et al*, 2009), while the HRDE-1 argonaute carries heritable endo-siRNAs that silence foreign, dangerous, or aberrant elements, whose expression could be deleterious, such as transposons (Luteijn *et al*, 2012; Shirayama *et al*, 2012; Rechavi, 2014). A strong and significant enrichment (**Figure 4C**) was found for endo-siRNAs which are carried in the germline by the argonautes HRDE-1 and WAGO-1 (Gu *et al*, 2009). These argonautes were found to be involved in gene silencing (Buckley *et al*, 2012; Gu *et al*, 2009), and HRDE-1 is required for inheritance of exogenous siRNAs (Buckley *et al*, 2012). A significant enrichment was also found for Mutator pathway small RNAs (Zhang *et al*, 2011), and putative piRNA targeted genes (fold change = 9.34, p-value < 0.0001(Bagijn *et al*, 2012)). On the contrary, a significant *depletion* was found for genes known to be targeted by CSR-1-carried small RNAs, a pathway that was suggested to support the expression of targeted genes (Claycomb *et al*, 2009; Shen *et al*, 2018). The helicase EMB-4 (Akay *et al*, 2017; Tyc *et al*, 2017) was shown to preferably bind introns of genes targeted by CSR-1; We could not detect a significant enrichment or depletion for genes whose introns are bound by EMB-4. All together, these results suggest that SET-25 is required for the maintenance of a specific sub-class of HRDE-1 and WAGO-1 small RNAs, associated with the Mutator and piRNA pathways, which target protein-coding (**Figure S2**).

### SET-25-dependent endo-siRNAs target a unique subset of newly evolved genes

What distinguishes the target genes of these SET-25-dependent endo-siRNAs? It was recently found that periodic A/T (PATC) sequences can shield germline genes from piRNA-induced silencing and allow germline expression of genes in H3K9me3-rich genomic regions (Frøkjær-Jensen *et al*, 2016; Zhang *et al*, 2018). Fittingly, we found that SET-25-dependent endo-siRNAs target genes that had significantly lower PATC density **(Figure 5A).** However, this feature is unlikely to distinguish between *oma-1* and *gfp*, since the *oma-1* gene has a very low PATC density (**Figure S3)**.

**Figure 5.**
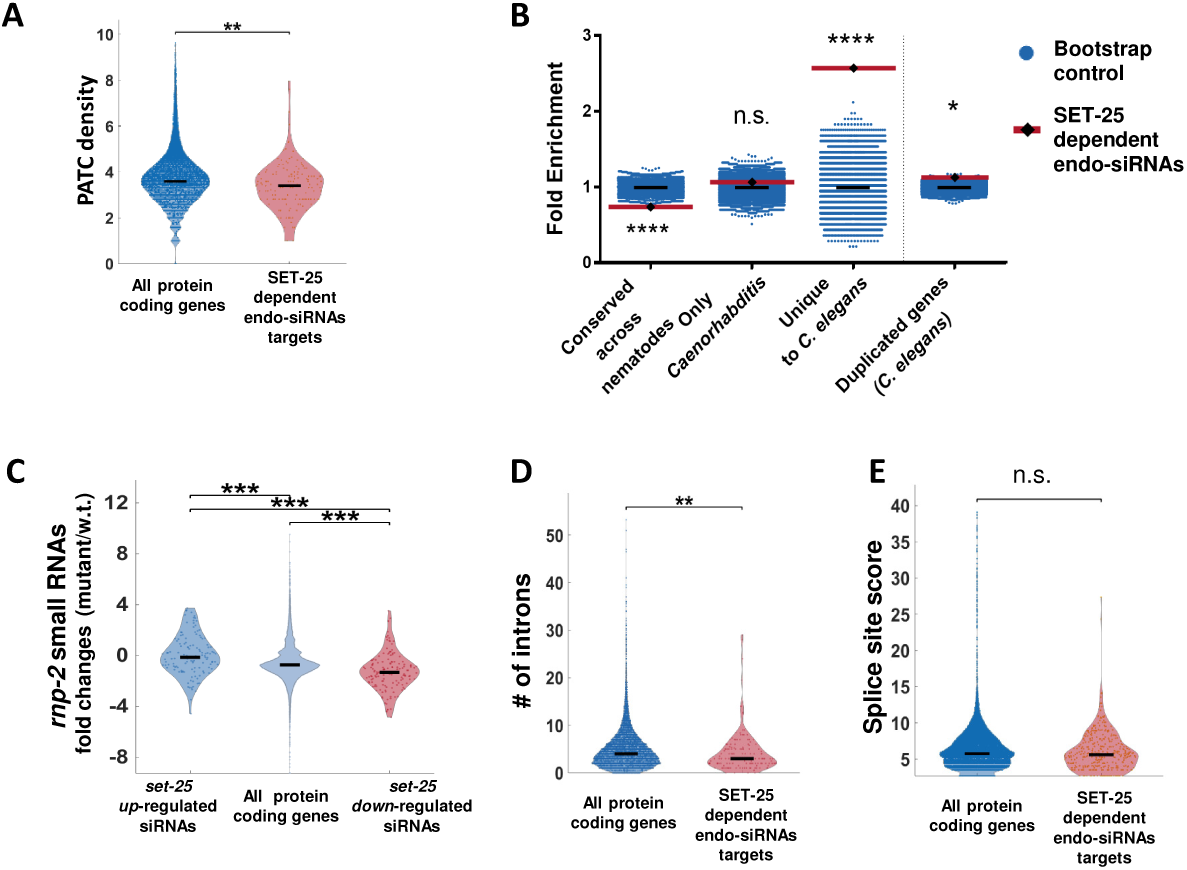
SET-25 dependent endo-siRNAs target newly evolved genes. (**A**) A PATC density analysis for SET-25 dependent endo-siRNAs gene targets. The PATC density values (obtained from (Frøkjær-Jensen *et al*, 2016)) for all protein-coding genes and SET-25 dependent endo-siRNAs gene targets are presented. p < 0.01, Wilcoxon rank sum test. For clarity of display, values are shown in log2 scale (after addition of 1). **(B)** An enrichment analysis of genes conserved at different levels, and duplicated genes amongst gene targets of SET-25 dependent endo-siRNAs. The gene sets were generated based on the homology field in WormBase that details the orthologs and paralogs of each nematode gene. We defined a duplicated gene as a gene that has a paralog in *C. elegans.* An enrichment value was calculated for each given gene set and control enrichment values were obtained from random gene sets with a comparable size. * p-value < 0.05 for duplicated gene set analysis, * p-value < 0.0001 for gene conservation enrichment analysis. (**C**) An analysis of endo-siRNAs fold changes in *rnp-2* mutants for genes targets of endo-siRNAs downregulated or upregulated in *set-*25 mutants or all genes. All p-values < 0.001, Wilcoxon rank sum test. (**D**) An analysis of intron numbers of gene targets of SET-25 dependent endo-siRNAs (obtained from (Lev *et al*, 2017)) compared to all protein-coding genes. In cases of genes that have more than one transcript, the average intron value is used.* p-value = 0.0053, Wilcoxon rank sum test. (**E**) An analysis of splicing motif divergence score (obtained from (Newman *et al*, 2018)) of gene targets of SET-25 dependent endo-siRNAs and all protein-coding genes. p-value > 0.05, Wilcoxon rank sum test..

The list of genes which are targeted by SET-25-dependnt endo-siRNAs was enriched for genes targeted by ERGO-1-dependent endo-siRNAs (fold change = 14.06, p-value < 0.0001, **Figure 4C**). Many of the genes which are targeted by ERGO-1-bound endo-siRNAs are duplicated genes (Vasale *et al*, 2010; Fischer *et al*, 2011). Still, only a modest (yet significant) enrichment was found for duplicated genes amongst the genes that had reduced endo-siRNA levels targeting them in *set-25* mutants (fold change = 1.12, p-value = 0.012 **Figure 5B**). An additional characteristic of the set of genes targeted by ERGO-1 endo-siRNAs is an enrichment for poorly conserved genes, that have fewer introns, and possess splicing site sequences that diverge from the consensus sequence (Newman *et al*, 2018; Fischer *et al*, 2011). It was recently suggested that these poorly conserved genes are targeted for silencing because their aberrant or “non-self-like” splicing signals are detected by the splicing machinery (Newman *et al*, 2018). Therefore, we examined whether SET-25-dependent endo-siRNA targets can be distinguished by their splicing signals.

The changes in the endo-siRNA pool in mutants of small nuclear ribonucleoprotein-associated protein RNP-2/U1A (*rnp-2*) mirrored the endo-siRNA changes found in *set-25* mutants (**Figure 5C**). We also found that genes targeted by SET-25-dependent endo-siRNAs bear fewer introns (**Figure 5D,** p = 0.0053), and were enriched with genes in which the introns are targeted by endo-siRNAs (**Figure S4A,** In most cases endo-siRNAs target only exons). Importantly, no significant differences in the length of the coding sequences were found, hence, the difference in intron number does not simply derive from differences in gene lengths (**Figure S4B**, p = 0.8673). We did not however, find significant differences in the splicing motif divergence score (obtained from (Newman *et al*, 2018)). Since splicing also directly affects the RNAi machinery untangling its role in endogenous RNAi is challenging (Newman *et al*, 2018). Thus, splicing may contribute to distinguishing genes targeted by SET-25 dependent endo-siRNAs.

Intriguingly, a significant enrichment for *newly evolved genes*, defined here as genes which had no orthologs outside *C. elegans*, was found among the SET-25-dependent endo-siRNA target genes (fold change = 2.57, 35/151 genes, p-value < 0.0001, **Figure 5B and Table S1**). Concordantly, in the same set we also found a significant *depletion* for nematode-conserved genes (fold change = 0.73, p-value < 0.0001, **Figure 5B**). In general, we find that certain endo-siRNA sub-classes, such as ERGO-1 and HRDE-1 bound small RNAs, exhibit a general enrichment for newly evolved genes (**Figure S5**). The significant enrichment for newly evolved genes among SET-25-dependent endo-siRNAs is maintained, however, even after excluding SET-25-endo-siRNA target genes that are also targeted by HRDE-1, ERGO-1, or WAGO-1 or Mutator endo-siRNAs (59 out of 151 genes are not shared, fold enrichment = 2.97, p-value = 0.0001). Further, we find that newly evolved genes have higher levels of H3K9me3 (**Figure S6**), Likewise, in the absence of RNAi, in wild-type animals, *gfp*, the newly evolved gene that we investigated, has higher levels of H3K9me3, in comparison to the well-conserved *oma-1* gene (**Figure 2B**). The fact that across the genome SET-25-dependent endo-siRNAs target newly evolved and H3K9me3 methylated genes (**Figure 4B** and **Figure 5B**), may explain why inheritance of RNAi responses raised against *gfp*, but not *oma-1*, depends on SET-25 (**Figure 1)**.

In summary, our experiments reveal a specific role for histone modifications in small RNA inheritance. While in S. *pombe* and *A. thaliana* a feedback between H3K9me3 and small RNAs was suggested to be required for silencing, the worm’s RNAi inheritance machinery may use H3K9me3 as a mark that distinguishes genes identified as “new”. Since newly evolved genes can be disruptive, small RNAs survey these H3K9me3-flagged elements transgenerationally,

## Discussion

Our study began from an investigation of a perplexing asymmetry in the requirement of specific H3K9 methyltransferases for heritable silencing of the endogenous gene *oma-1* and the foreign gene *gfp*. Single mutants of *set-25* and *set-32* and the *met-2;set-25;set-32* triple mutant displayed different heritable dynamics when either the *gfp* or the *oma-1* gene were targeted by RNAi. These results are not unique to the specific *gfp* transgene that was tested, since similar observations have been made with other transgenes (Shirayama *et al*, 2012; Klosin *et al*, 2017; Spracklin *et al*, 2017; Lev *et al*, 2017).

Unlike mutations in these histone methyltransferases, which negatively affect heritable silencing of *gfp*, but not *oma-1*, mutations in genes required for small RNA inheritance negatively affect heritable silencing of both *oma-1* and *gfp.* For example, the argonaute HRDE-1 is required for inheritance of RNAi responses against both targets (Buckley *et al*, 2012; Weiser *et al*, 2017; Ashe *et al*, 2012; Shirayama *et al*, 2012; Kalinava *et al*, 2017). The fact that heritable RNAi responses aimed at different genes are affected by different proteins should be taken into account when studying transgenerational inheritance. Specifically, when screening for genes that affect such inheritance, one must acknowledge that heritable silencing of different targets requires different chromatin modifiers.

Future studies will hopefully reveal why some recently evolved genes, but not others, display high levels of H3K9me3 (in the absence of RNAi), and are targeted by endo-siRNAs. Recent studies examined why transgenes are sensitive to silencing by synthetic piRNAs, while endogenous germline expressed genes, including *oma-1*, are not. This protection was suggested to be conferred at least in part by PATC sequences, and to be independent of the genomic location of the gene (Zhang *et al*, 2018). PATC sequences were previously shown to allow expression of transgenes in the germline in heterochromatic areas (Frøkjær-Jensen *et al*, 2016). Similarly, our analysis revealed that the gene targets of SET-25-dependent endo-siRNAs have lower levels of PATC density (**Figure 5A**). However, the *oma-1* gene does not possess many PATC sequences (**Figure S3**). An additional theory suggested that an intrinsic unknown coding-sequence feature confers resistance to silencing by piRNAs. Seth et al. have studied why a fusion between *oma-1* and *gfp* can trans-activate silenced *gfp* transgenes (an effect known as “RNAa”,(Seth *et al*, 2013)). While unique “protecting” sequence features were not described in that work, the authors showed that an unknown coding-sequence feature, not related to the codon usage or the translation of the protein, grants the *oma-1* gene with its ability to activate silenced transgenes (Seth *et al*, 2018). It is possible that the gene targets of SET-25-dependent small RNAs that we describe here have unique intrinsic sequences that distinguish them as well. The different requirement of methyltransferases for heritable silencing of some genes but not others may be related to such intrinsic sequence features. Alternatively, it is possible, as was suggested in the past, that new genes are silenced because they are not licensed transgenerationally by heritable small RNAs for expression (Claycomb *et al*, 2009; Shen *et al*, 2018). If this is the case, future studies will hopefully reveal how such license is granted (See **Figure 6** for Scheme).

**Figure 6.**
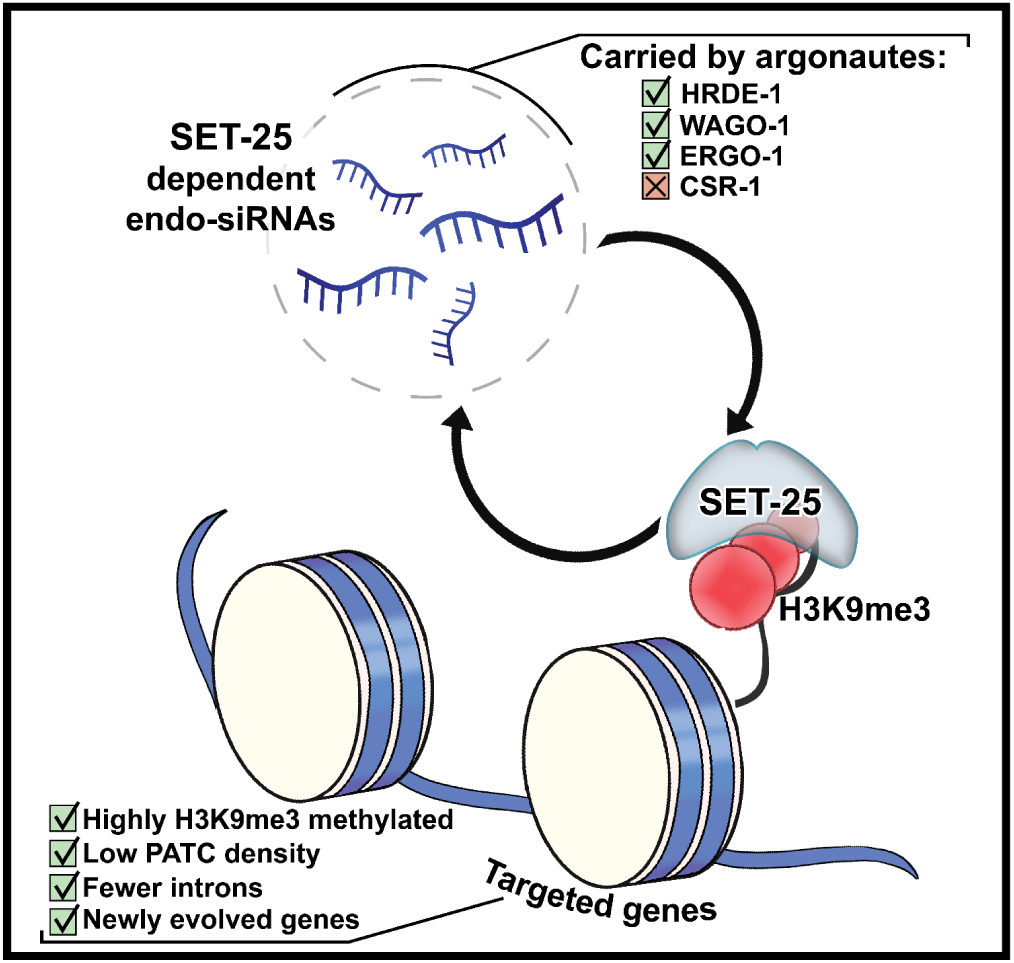
Scheme characterizing SET-25 dependent endo-siRNAs and their targets. SET-25 dependent endo-siRNAs are enriched with small RNAs known to be carried by the argonautes HRDE-1, WAGO-1 and ERGO-1 but not CSR-1. SET-25 dependent endo-siRNAs targets are enriched with newly evolved genes, genes bearing fewer introns, fewer PATC sequences, and are marked with higher levels of H3K9me3.

## Materials and Methods

### Cultivation of the Worms

Standard culture techniques were used to maintain the nematodes on nematode growth medium (NGM) plates seeded with OP50 bacteria. Extreme care was taken to avoid contamination or starvation, and contaminated plates were discarded from the analysis.

### Strains used in this study

BFF24 [*set-32(ok1457)* I.; SX1263 mjIs134 II. [p*mex-5::gfp::h2b::tbb-2* II.]

BFF25 [*set-32(ok1457)* I.; *oma-1(zu405)* IV.]

BFF26 [*met-2(n4256)* III.; *set-25(n5021)* III.; *set-32(ok1457)* I.; SX1263 *mjIs134* II. *[pmex-5::gfp::h2b::tbb-2* II.]

BFF27 [*met-2(n4256)* III.; *set-25(n5021)* III.; *set-32(ok1457)* I.; *oma-1(zu405)* IV.]]

BFF28 [*met-2(n4256)* III.; *set-32(ok1457)* I.; SX1263 mjIs134 II. [p*mex-5::gfp::h2b::tbb-2* II.]]

### RNAi bacteria

HT115 *Escherichia coli* strains expressing dsRNAs were used: anti-*oma-*1 RNAi bacteria were obtained from the Ahringer RNAi library (Kamath & Ahringer, 2003). For the sequence of the anti-*gfp* RNAi see supplemental data.

### RNAi experiments

RNAi HT115 *E.coli* bacteria were incubated in Lysogeny broth (LB) containing Carbenicillin (25 μg/mL) at 37°C overnight with shaking. Bacterial cultures were seeded onto NGM plates containing isopropyl β-D-1-thiogalactopyranoside (IPTG; 1 mM) and Carbenicillin (25 μg/mL) and grown overnight in the dark at room temperature. Five L4 animals were placed on RNAi bacteria plates and control empty-vector bearing HT115 bacteria plates and maintained at 20°C for 2 days and then removed. The progeny hatching on these plates was termed the P0 generation. In the next generations the worms were grown on *E.coli* OP50 bacteria. For anti-*gfp* RNAi experiments, four L4 animals were placed on plates for two days to lay the next generation. In every generation approximately 60 one day adult worms were collected and photographed per condition (see below). For anti-*oma-1* RNAi experiments, in each generation twelve individual L4 staged worms were placed in individual wells of a twelve well plate. Four days later the number of fertile worms was assessed (at least one progeny) and twelve individual L4 progeny worms were chosen from the most fertile well to continue to the next generation.

### Germline GFP expression analysis

Percentage silencing analysis: for each condition, around 60 animals were mounted on 2% agarose slides and paralyzed in a drop of M9 with 0.01% levamisole/0.1% tricaine. The worms were photographed with 10x objective using a BX63 Olympus microscope (Exposure time of 200 ms, and gain of 2). The images were analyzed with ImageJ2 software, and the percentage of worms lacking any germline GFP signal was calculated. GFP expression level analysis: for each condition, the GFP fluorescence level of the background and of oocyte nuclei of at least 30 worms was calculated using ImageJ2.

CTCF value was calculated as follows: CTCF = Integrated density of selected object X – (area of selected object X * mean fluorescence of background readings). The obtained CTCF value was normalized to the average CTCF value obtained from photographs of control animals of the same genotype, generation and age which were fed on control plates.

### Chromatin immunoprecipitation

Chromatin immunoprecipitation experiments were conducted as described in (Lev *et al*, 2017). For anti-H3K9me3 ChIP experiments the abcam, ab8898 antibodies were used.

### qPCR reactions

All Real time PCR reactions were performed using the KAPA SYBR Fast qPCR and run in the Applied Biosystems 7300 Real Time PCR System.

The primer sequences used in qRT-PCR:

*gfp* set #1 FOR: ACACAACATTGAAGATGGAAGC

*gfp* set #1 REV: GACAGGTAATGGTTGTCTGG

*gfp* set #2 FOR: GTGAGAGTAGTGACAAGTGTTG

*gfp* set #2 REV: CTGGAAAACTACCTGTTCCATG

*oma-1* set#1 FOR: AACTTTGCCCGTTTCACC

*oma-1* set#1 REV: TCAAGTTAGCAGTTTGAGTAACC

*oma-1* set#2 FOR: TTGTTAAGCATTCCCTGCAC

*oma-1* set#2 REV: TCGATCTTCTCGTTGTTTTCA

(The above primer set was adapted from (Spracklin *et al*, 2017))

*dpy-28* FOR: CTGATGGATCCAGAGTTGG

*dpy-28* REV: CTGCTATACGCATCCTGTTC

*eft-3* FOR: CCAACATGATTAGTCAGATGACC

*eft-3* REV: CTAGGAGTTAGATGTGCAGG.

### Bioinformatic genome-wide endo-siRNAs analysis

Small RNA analysis were conducted as previously described (Lev *et al*, 2017). Briefly, Adapters were cut from the reads using Cutadapt (Martin, 2011). Reads that were not cut or were less than 19bp long, were removed. The quality of the libraries was assessed by FastQC (http://www.bioinformatics.babraham.ac.uk/projects/fastqc/). Reads were mapped to the *C. elegans* genome (WS235) using Bowtie2 (Langmead & Salzberg, 2012). The mapped reads were then counted using the python script HTseq_count (Langmead & Salzberg, 2012) using. gff feature file from wormbase.org (version WBcel235). Differential expression was analyzed using DESeq2 (Love *et al*, 2014). p-adjusted value < 0.1 was regarded as statistically significant. GEO accession GSE94798.

### Bioinformatic gene enrichment analysis

The enrichment values denote the ratio between (A) the observed representations of a specific gene set within a defined differentially expressed genes group, to (B) the expected one, i.e., the representation of the examined gene set among all protein-coding genes in *C. elegans*. The analysis was done for 8 gene sets: (1) 7727 genes enriched in oocytes gonads (Ortiz *et al*, 2014) and 9012 genes enriched in spermatogenic gonads (Ortiz *et al*, 2014); we excluded genes with expression lower than 1 (2) 11427 genes expressed in isolated neurons (Kaletsky *et al*, 2015). (3) 7176 genes expressed in intestine (Gerstein *et al*, 2010) (4) 2957 genes expressed in pharynx (Gerstein *et al*, 2010) (5) 2526 genes expressed in body muscle (Gerstein *et al*, 2010) (6) 4146 targets of CSR-1 (Claycomb *et al*, 2009) (7) 1478 targets of HRDE-1 (Buckley *et al*, 2012) (8) 87 targets of WAGO-1 (Gu *et al*, 2009) (9) 399 targets of ALG-3/4 class small RNAs (Conine *et al*, 2010) (10) 1823 targets of mutator class small RNAs (11) 721 EGO-1 dependent small RNA gene targets (Maniar & Fire, 2011), (12) 23 gene targets of small RNAs up-regulated in *ego-1* mutants (Maniar & Fire, 2011), (13) 49 genes targeted by 26G-RNAs enriched in ERGO-IP (Vasale *et al*, 2010) (14) 77 genes depleted of 22G-RNAs in *ergo-1* mutants (Vasale et al. 2010), and (15) 348 putative piRNA gene targets (Bagijn *et al*, 2012). The putative piRNA gene targets were defined as genes for which, in at least one transcript, the ratio of the # 22G-RNA reads at piRNA target sites between wild type to *prg-1* is at least 2 (linear scale). Note that the indicated number above achieved after intersection between the various published data sources and the records appears in the *.gff file used by us.

The enrichment value of a given gene set i in differentially expressed gene targeting small RNAs was calculated using the following formula:

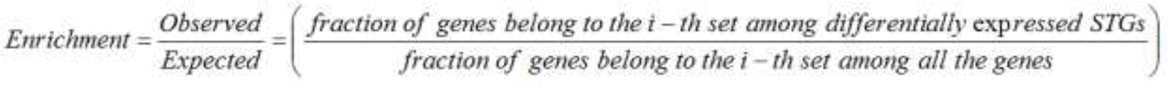

Obtaining the observed-to-expected ratios, we then calculated the corresponding p-values using 10,000 random gene groups identical in size to that of the examined group of differentially expressed genes.

### Bioinformatic H3K9me3 signal analysis

H3K9me3 signals are based on (McMurchy *et al*, 2017), the shown signal represents the averaged H3K9me3 signal in two replicates of young adults (GEO accession GSE87524).

### Gene sets by conservation

The classification of gene sets by conservation was done by mining the “Homology” field of all the *C. elegans* protein-coding genes in WormBase (www.wormbase.com). We defined the following three gene sets (Figure 5B):

1. Unique to *C. elegans* – *C. elegans* genes which have no orthologues gene in any of the following species: *B. malayi, C. brenneri, C. briggsae, C. japonica, C. remanei, O. volvulus, P. pacificus* and *S. ratti*.
2. *Caenorhabditis* only - *C. elegans* genes which have at least one orthologues gene in one of the *C. brenneri, C. briggsae, C. remanei* and *C. japonica* species, and have no orthologues gene in any of the *B. malayi, O. volvulus, P. pacificus* and *S. ratti* species.
3. Conserved among nematodes - *C. elegans* genes which have at least one orthologues gene in one of the *C. brenneri, C. briggsae, C. remanei* and *C. japonica* species, and in addition have at least one orthologues gene in one of the *B. malayi, O. volvulus, P. pacificus* and *S. ratti* species.

### Statistical analysis

For RNAi experiments, Two-way ANOVA tests were used to compare the percentages of the RNAi-affected worms (GFP silencing or fertility for the oma-1 assay) between the tested genotypes. In cases of multiple comparisons between genotypes and across generations, Sidak multiple comparison tests were applied. For GFP fluorescence experiments, Two-way ANOVA tests were used to compare the normalized GFP expression levels between the genotypes and across the biological repeats. For H3K9me3 qPCR-ChIP experiments Two-way ANOVA tests were used to compare the delta-delta-Ct (or delta-Ct) values between the gfp and the oma-1 loci obtained using two different primer sets. In cases of comparisons between genotypes and loci the Sidak multiple comparison tests were applied. Biological replicates were performed using separate populations of animals. Statistical tests were performed using GraphPad Prism software (Graphpad Prism) version 6. The statistical analysis used for each of the bioinformatics analyses is listed under the corresponding bioinformatics methods.

## Acknowledgements

We thank all the Rechavi lab members for the helpful comments and fruitful discussions. Some strains were provided by the CGC, which is funded by NIH Office of Research Infrastructure Programs (P40 OD010440). We thank Yosef Shiloh, Yael Ziv, for their assistance and advice. Special thanks to Dror Cohen for the illustrations that he contributed. This work was supported by the ERC (grant #335624) and the Israel Science Foundation (grant #1339/17) and. O.R. gratefully acknowledges the support of the Allen Discovery Center of the Paul G. Allen Frontiers Group.

## Author contributions

**I.L.** and **O.R.** conceived and designed the experiments; **I.L.** performed the experiments; **H.G.** performed bioinformatics data analysis; and **I.L.**, **H.G.** and **O.R.** wrote the paper.

## Conflict of interest

None.

## Conventions and Abbreviations

RNAi: RNA interference
Endo-siRNAs: Endogenous small interfering RNAs
ChIP: Chromatin Immunoprecipitation
qPCR: Quantitative real time PCR
PATC: periodic A/T (PATC) sequences

## Declarations

None.

## Supplemental Figure Legends

**Figure S1.**
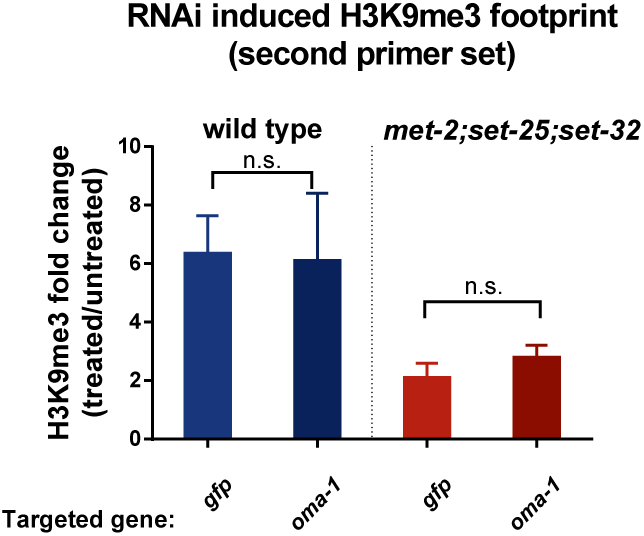
Analysis of protein-coding genes with significantly increased endo-siRNAs levels targeting them in *set-25* mutants compared to wild type. (**A**) An analysis of H3K9me3 signals (based on published data from (McMurchy *et al*, 2017)) on gene targets of endo-siRNAs *up-*regulated in *set-25* mutants (based on (Lev *et al*, 2017), Blue). All genes are aligned according to their Transcription Start Sites (TSS), and the regions of 1000 base pairs upstream and downstream of the TSS are shown on the x axis. The y axis shows the averaged signal of the H3K9me3 modification as a function of distance from the TSS (**B**) An enrichment analysis for expression in specific tissues (left) and an enrichment analysis for different small RNA pathways (right) are presented. Fold enrichment values are color coded. p-values were calculated using 10,000 random gene sets identical in their size to the examined gene set. Asterisk denotes statistically significant enrichment values (p-value < 0.05).

**Figure S2.**
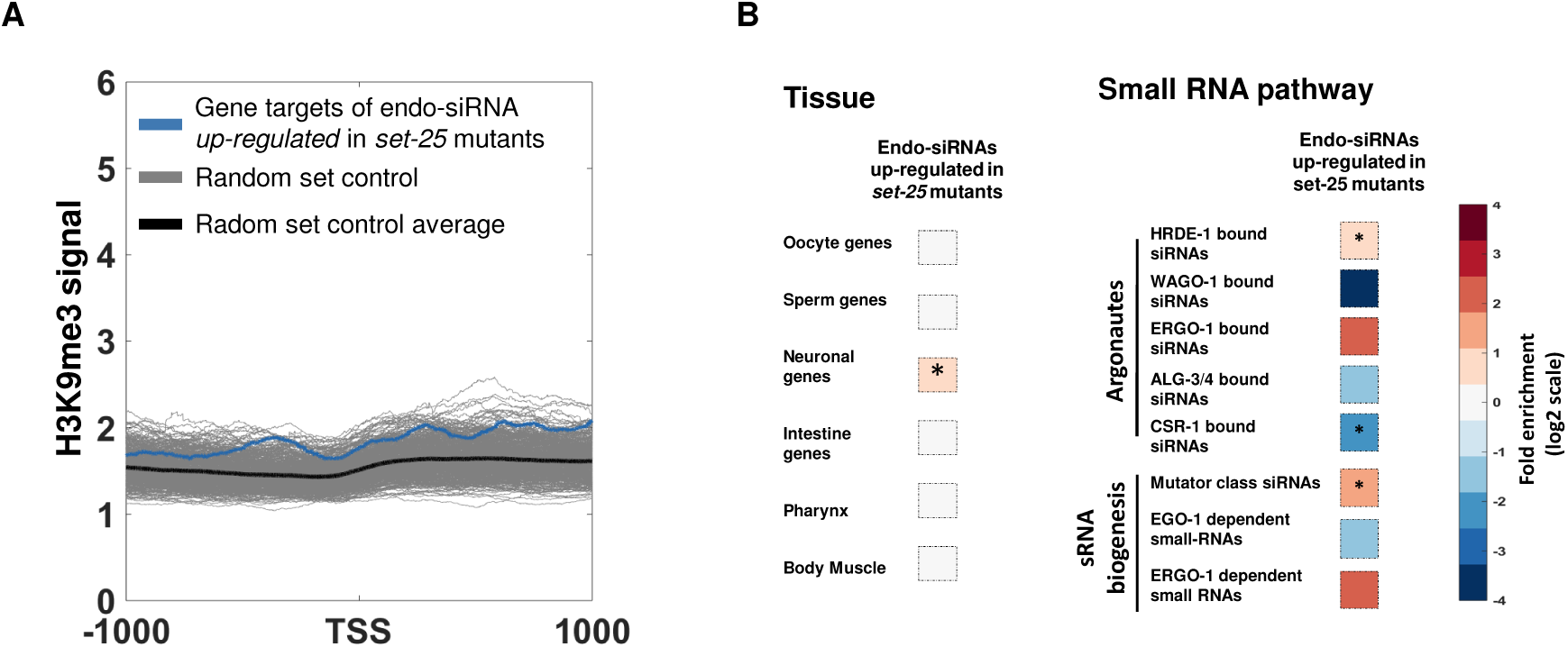
Unique and shared genes targets of SET-25 dependent endo-siRNAs. A Venn diagram analysis of shared gene targets of HRDE-1 (Buckley *et al*, 2012), Mutator pathway (Phillips *et al*, 2014), ERGO-1 dependent small RNA targets (Vasale *et al*, 2010) and SET-25 (Lev *et al*, 2017) dependent small RNAs.

**Figure S3.**
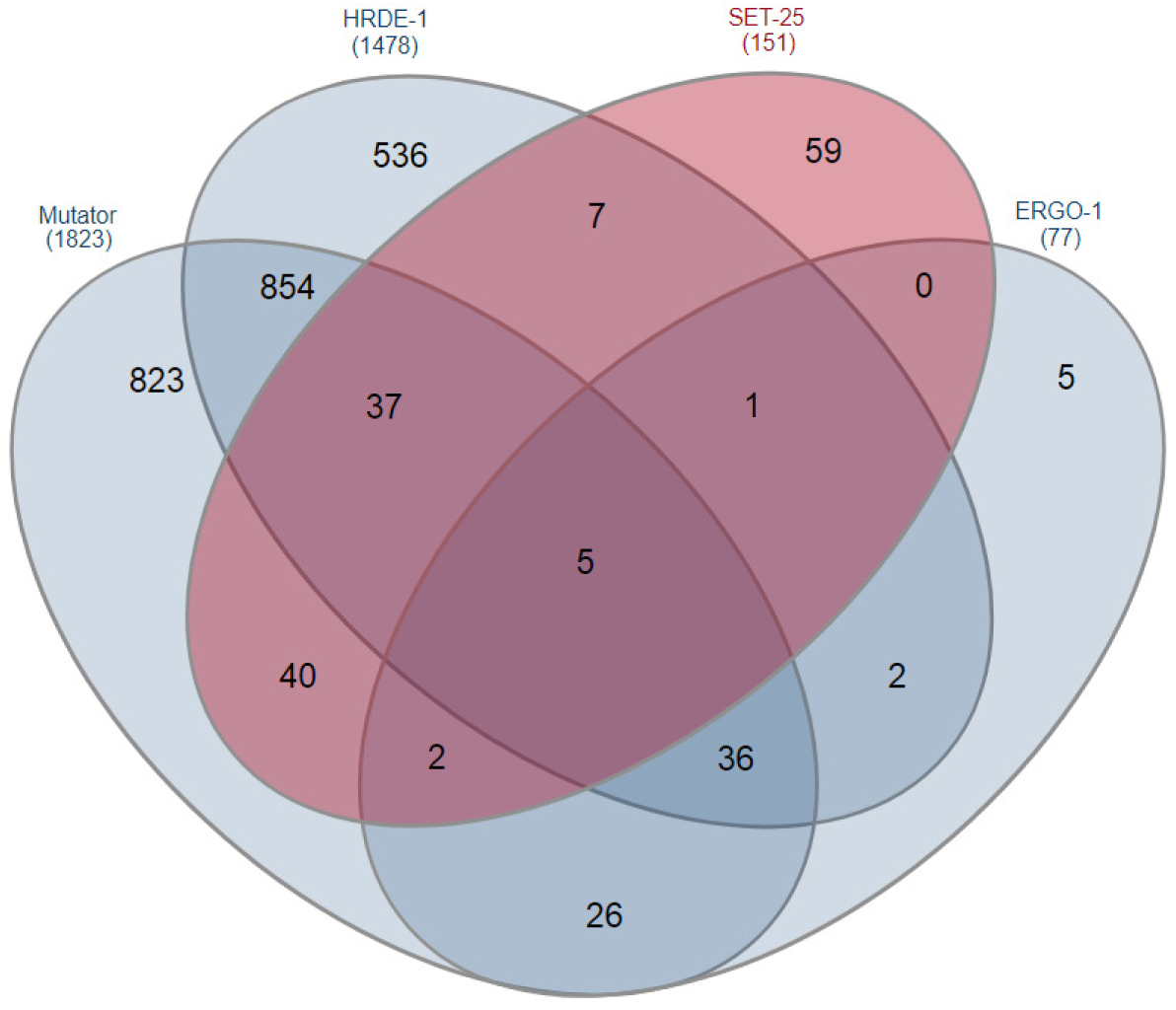
The *oma-1* gene bears low density of PATC. A histogram of PATC density of all of *C. elegans* genes (Blue) and the PATC density of the *oma-1* gene (Red). The median PATC density of all genes=11.00 (marked with black vertical line), 243 points counted in the rightmost bin have values greater than 178.47.

**Figure S4.**
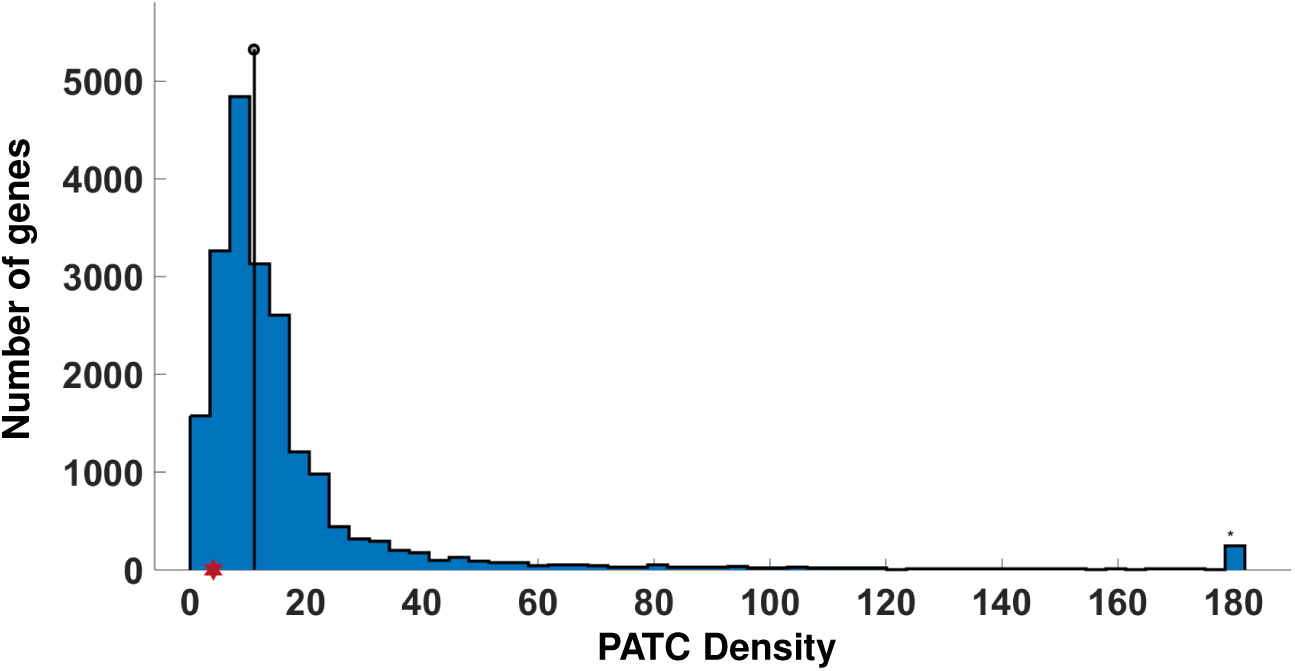
Genes targeted by SET-25 dependent endo-siRNAs gene targets are more targeted by small RNAs on introns compared to other genes. **(A)** The proportion of genes with different ratios of small RNAs aligned to Intron-Exon-junction / Exon-Exon junction (obtained from (Newman *et al*, 2018)) are presented. The control proportion values obtained from 10,000 random gene sets with an identical size to SET-25 dependent end-siRNA gene targets set are shown in grey. The proportion value of SET-25 dependent endo-siRNA gene targets is shown in red. (**B**) The lower intron numbers found in genes targeted by SET-25 dependent endo-siRNAs are not explained by smaller coding sequence sizes. An analysis of coding sequence length of genes targeted by SET-25 dependent endo-siRNAs and all protein-coding genes. p-value = 0.8673, Wilcoxon rank sum test. This figure relates to **Figure 5D.**

**Figure S5.**
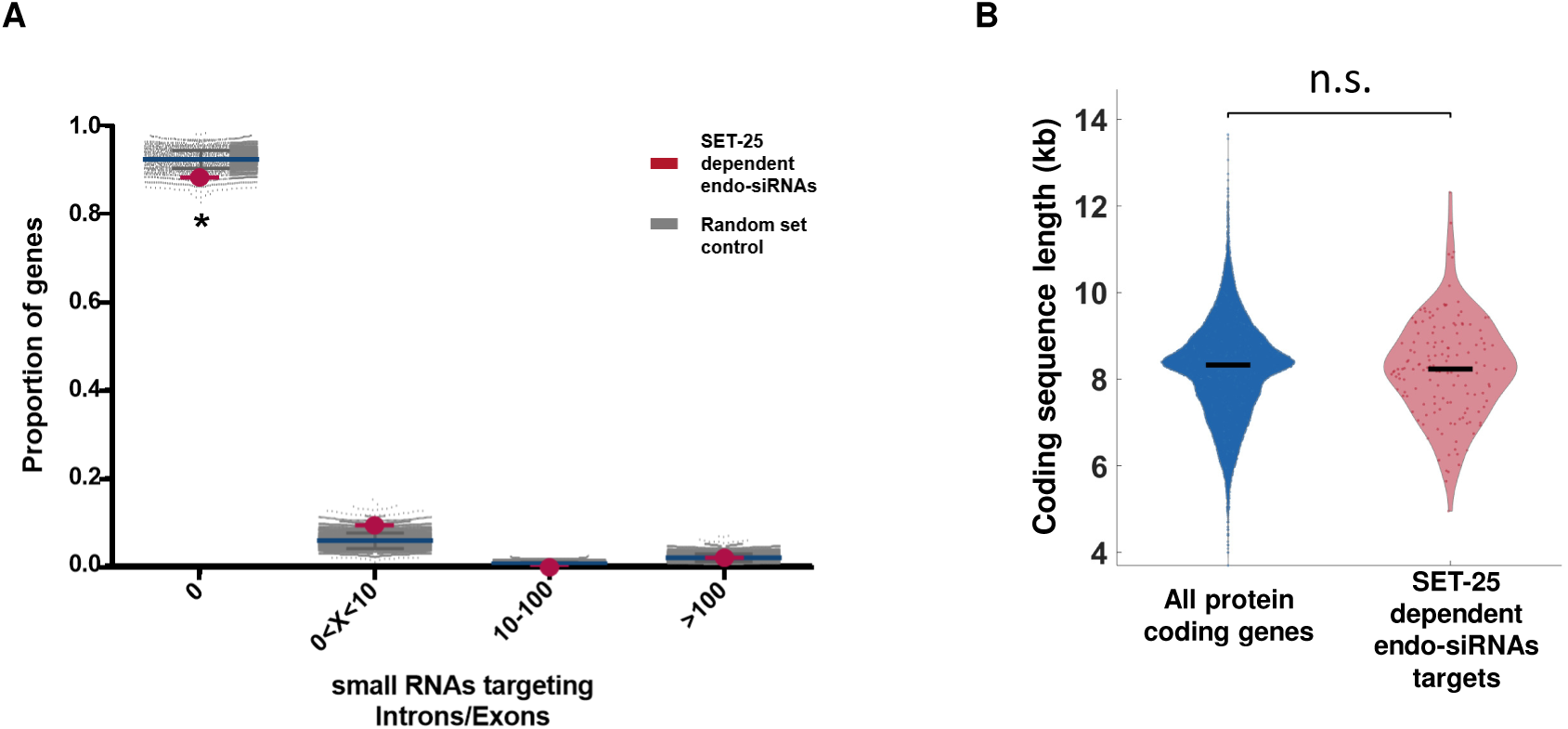
An enrichment analysis of newly acquired genes amongst genes targeted by known endo-siRNA pathways. An enrichment analysis for newly acquired genes, bearing no orthologues among other nematodes amongst different endo-siRNAs pathways (the newly acquired gene set was generated based on the “Homology” field of the *C. elegans* genes in WormBase). Fold enrichment values are color coded. Asterisk denotes statistically significant enrichment values (p-value < 0.001). P-values were calculated using 10,000 random gene sets identical in their size to the examined gene set.

**Figure S6.**
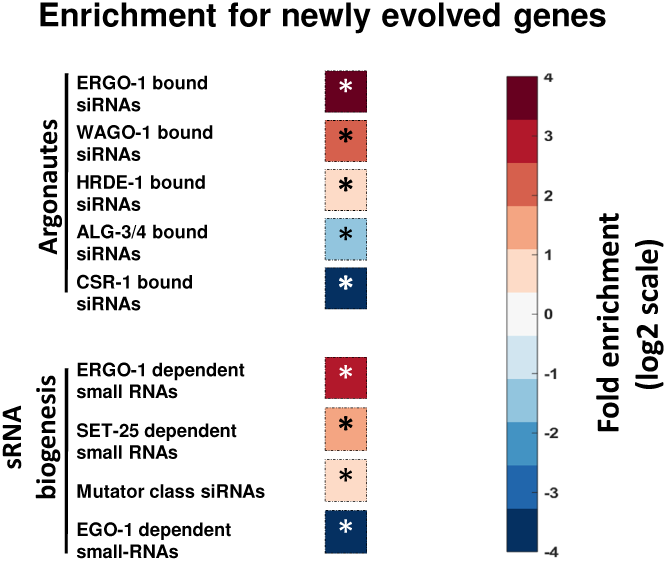
Newly acquired genes exhibit enhanced levels of H3K9me3. Analysis of H3K9me3 levels (based on published data from (McMurchy *et al*, 2017)) of newly acquired genes in *C. elegans.* As a control, shown are the H3K9me3 signature of 500 random gene sets identical in size to the gene set of interest.

**Figure S7.**
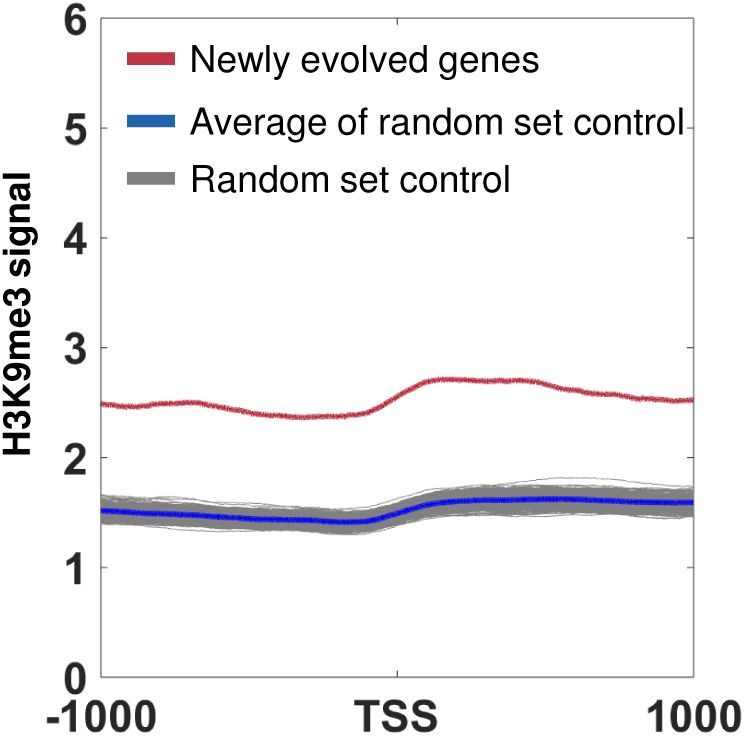

**Supplemental Table 1 – list of SET-25-dependent endo-siRNAs targets and their conservation.** The table includes all the *C. elegans* genes we examined (20,447 genes). Each gene is marked for being: A. a target of SET-25 dependent endo-siRNAs, B. Conserved among nematodes, C. Conserved only in *Caenorhabditis*, D. Unique to *C. elegans.*

**Supplemental File 1 – Sequence of anti-*gfp* RNAi targeting dsRNA used in this study.**

## References and Citations

Akay A, Di Domenico T, Suen KM, Nabih A, Parada GE, Larance M, Medhi R, Berkyurek AC, Zhang X, Wedeles CJ, Rudolph KLM, Engelhardt J, Hemberg M, Ma P, Lamond AI, Claycomb JM & Miska EA (2017) The Helicase Aquarius/EMB-4 Is Required to Overcome Intronic Barriers to Allow Nuclear RNAi Pathways to Heritably Silence Transcription. Dev. Cell 42: 241–255.e6

Alcazar RM,Lin R & Fire AZ (2008) Transmission Dynamics of Heritable Silencing Induced by Double-Stranded RNA in Caenorhabditis elegans. 1288: 1275–1288

Ashe A, Sapetschnig A, Weick EM, Mitchell J, Bagijn MP, Cording AC, Doebley AL, Goldstein LD, Lehrbach NJ, Le Pen J, Pintacuda G, Sakaguchi A, Sarkies P, Ahmed S & Miska EA (2012) PiRNAs can trigger a multigenerational epigenetic memory in the germline of C. elegans. Cell 150: 88–99

Bagijn MP, Goldstein LD, Sapetschnig A, Weick E-M, Bouasker S, Lehrbach NJ, Simard MJ & Miska EA (2012) Function, Targets, and Evolution of Caenorhabditis elegans piRNAs. Science (80-.). 337:

Bessler JB, Andersen EC & Villeneuve AM (2010) Differential localization and independent acquisition of the H3K9me2 and H3K9me3 chromatin modifications in the Caenorhabditis elegans adult germ line. PLoS Genet. 6: e1000830

Buckley B a, Burkhart KB, Gu SG, Spracklin G, Kershner A, Fritz H, Kimble J, Fire A & Kennedy S (2012) A nuclear Argonaute promotes multigenerational epigenetic inheritance and germline immortality. Nature 489: 447–51

Castel SE & Martienssen RA (2013) RNA interference in the nucleus: roles for small RNAs in transcription, epigenetics and beyond. Nat. Rev. Genet. 14: 100–12

Chicas A, Forrest EC, Sepich S, Cogoni C & Macino G (2005) Small Interfering RNAs That Trigger Posttranscriptional Gene Silencing Are Not Required for the Histone H3 Lys9 Methylation Necessary for Transgenic Tandem Repeat Stabilization in Neurospora crassa. Mol. Cell. Biol. 25: 3793–3801

Claycomb JM, Batista PJ, Pang KM, Gu W, Vasale JJ, van Wolfswinkel JC, Chaves DA, Shirayama M, Mitani S, Ketting RF, Conte D & Mello CC (2009) The Argonaute CSR-1 and Its 22G-RNA Cofactors Are Required for Holocentric Chromosome Segregation. Cell 139: 123–134

Conine CC, Batista PJ, Gu W, Claycomb JM, Chaves DA, Shirayama M & Mello CC (2010) Argonautes ALG-3 and ALG-4 are required for spermatogenesis-specific 26G-RNAs and thermotolerant sperm in Caenorhabditis elegans. Proc. Natl. Acad. Sci. 107: 3588–93

Daniel Holoch, Danesh Moazed, Holoch D & Moazed D (2015) RNA-mediated epigenetic regulation of gene expression. Nat. Rev. Genet. 16: 71–84

Fischer SEJ, Montgomery TA, Zhang C, Fahlgren N, Breen PC, Hwang A, Sullivan CM, Carrington JC & Ruvkun G (2011) The ERI-6/7 Helicase Acts at the First Stage of an siRNA Amplification Pathway That Targets Recent Gene Duplications. PLoS Genet. 7: e1002369

Frøkjær-Jensen C, Jain N, Hansen L, Davis MW, Li Y, Zhao D, Rebora K, Millet JRM, Liu X, Kim SK, Dupuy D, Jorgensen EM & Fire AZ (2016) An Abundant Class of Non-coding DNA Can Prevent Stochastic Gene Silencing in the C. elegans Germline. Cell 166: 343–357

Gammon DB, Ishidate T, Li L, Gu W, Silverman N & Mello CC (2017) The Antiviral RNA Interference Response Provides Resistance to Lethal Arbovirus Infection and Vertical Transmission in Caenorhabditis elegans. Curr. Biol. 27: 795–806

Gerstein MB, Lu ZJ, Van Nostrand EL, Cheng C, Arshinoff BI, Liu T, Yip KY, Robilotto R, Rechtsteiner A, Ikegami K, Alves P, Chateigner A, Perry M, Morris M, Auerbach RK, Feng X, Leng J, Vielle A, Niu W, Rhrissorrakrai K, et al (2010) Integrative Analysis of the Caenorhabditis elegans Genome by the modENCODE Project. Science (80-.). 330: 1775–1787

Gu SG, Pak J, Guang S, Maniar JM, Kennedy S & Fire A (2012) Amplification of siRNA in Caenorhabditis elegans generates a transgenerational sequence-targeted histone H3 lysine 9 methylation footprint. Nat. Genet. 44: 157–64

Gu W, Shirayama M, Conte D, Vasale J, Batista PJ, Claycomb JM, Moresco JJ, Youngman EM, Keys J, Stoltz MJ, Chen C-CGCG, Chaves DA, Duan S, Kasschau KD, Fahlgren N, Yates JR, Mitani S, Carrington JC & Mello CC (2009) Distinct Argonaute-Mediated 22G-RNA Pathways Direct Genome Surveillance in the C. elegans Germline. Mol. Cell 36: 231–244

Guang S, Bochner AF, Burkhart KB, Burton N, Pavelec DM & Kennedy S (2010) Small regulatory RNAs inhibit RNA polymerase II during the elongation phase of transcription. Nature 465: 1097–1101

Kaletsky R, Lakhina V, Arey R, Williams A, Landis J, Ashraf J & Murphy CT (2015) The C. elegans adult neuronal IIS/FOXO transcriptome reveals adult phenotype regulators. Nature 529: 92–96

Kalinava N, Ni J, Gajic Z, Ushakov H & Gu S (2018) Caenorhabditis elegans heterochromatin factor SET-32 plays an essential role in transgenerational establishment of nuclear RNAi-mediated epigenetic silencing. bioRxiv: 255562

Kalinava N, Ni JZ, Peterman K, Chen E & Gu SG (2017) Decoupling the downstream effects of germline nuclear RNAi reveals that H3K9me3 is dispensable for heritable RNAi and the maintenance of endogenous siRNA-mediated transcriptional silencing in Caenorhabditis elegans. Epigenetics and Chromatin 10: 6

Kamath RS & Ahringer J (2003) Genome-wide RNAi screening in Caenorhabditis elegans. Methods 30: 313–321

Klosin A, Casas E, Hidalgo-Carcedo C, Vavouri T & Lehner B (2017) Transgenerational transmission of environmental information in C. elegans. Science (80-.). 356: 320–323

Langmead B & Salzberg SL (2012) Fast gapped-read alignment with Bowtie 2. Nat. Methods 9: 357–359

Lev I, Seroussi U, Gingold H, Bril R, Anava S & Rechavi O (2017) MET-2-Dependent H3K9 Methylation Suppresses Transgenerational Small RNA Inheritance. Curr. Biol. 27: 1138–1147

Love MI, Huber W & Anders S (2014) Moderated estimation of fold change and dispersion for RNA-seq data with DESeq2. Genome Biol. 15: 550

Luteijn MJ, van Bergeijk P, Kaaij LJT, Almeida MV, Roovers EF, Berezikov E & Ketting RF (2012) Extremely stable Piwi-induced gene silencing in Caenorhabditis elegans. EMBO J. 31: 3422–3430

Luteijn MJ & Ketting RF (2013) PIWI-interacting RNAs: from generation to transgenerational epigenetics. Nat. Rev. Genet. 14: 523–534

Malone CD & Hannon GJ (2009) Small RNAs as guardians of the genome. Cell 136: 656–68

Maniar JM & Fire AZ (2011) EGO-1, a C. elegans RdRP, modulates gene expression via production of mRNA-templated short antisense RNAs. Curr. Biol. 21: 449–459

Mao H, Zhu C, Zong D, Weng C, Yang X, Huang H, Liu D, Feng X & Guang S (2015) The Nrde Pathway Mediates Small-RNA-Directed Histone H3 Lysine 27 Trimethylation in Caenorhabditis elegans. Curr. Biol. 25: 2398–2403

Martin M (2011) Cutadapt removes adapter sequences from high-throughput sequencing reads. EMBnet.journal 17: 10–12

McMurchy AN, Stempor P, Gaarenstroom T, Wysolmerski B, Dong Y, Aussianikava D, Appert A, Huang N, Kolasinska-Zwierz P, Sapetschnig A, Miska EA & Ahringer J (2017) A team of heterochromatin factors collaborates with small RNA pathways to combat repetitive elements and germline stress. Elife 6: e21666

Minkina O & Hunter CP (2017) Stable Heritable Germline Silencing Directs Somatic Silencing at an Endogenous Locus. Mol. Cell 65: 659–670.e5

Moazed D, Bühler M, Buker SM, Colmenares SU, Gerace EL, Gerber SA, Hong E-JE, Motamedi MR, Verdel A, Villén J & Gygi SP (2006) Studies on the mechanism of RNAi-dependent heterochromatin assembly. Cold Spring Harb. Symp. Quant. Biol. 71: 461–71

Molnar A, Melnyk CW, Bassett A, Hardcastle TJ, Dunn R & Baulcombe DC (2010) Small silencing RNAs in plants are mobile and direct epigenetic modification in recipient cells. Science 328: 872–5

Newman MA, Ji F, Fischer SEJ, Anselmo A, Sadreyev RI & Ruvkun G (2018) The surveillance of pre-mRNA splicing is an early step in C. elegans RNAi of endogenous genes. Genes Dev. 32: 670–681

Ortiz MA, Noble D, Sorokin EP & Kimble J (2014) A new dataset of spermatogenic vs. oogenic transcriptomes in the nematode Caenorhabditis elegans. G3 (Bethesda). 4: 1765–72

Phillips CM, Montgomery BE, Breen PC, Roovers EF, Rim YS, Ohsumi TK, Newman MA, Van Wolfswinkel JC, Ketting RF, Ruvkun G & Montgomery TA (2014) MUT-14 and SMUT-1 DEAD box RNA helicases have overlapping roles in germline RNAi and endogenous siRNA formation. Curr. Biol. 24: 839–844

Rechavi O (2014) Guest list or black list: Heritable small RNAs as immunogenic memories. Trends Cell Biol. 24: 212–220

Rechavi O, Houri-Ze’evi L, Anava S, Goh WSS, Kerk SY, Hannon GJ & Hobert O (2014) Starvation-Induced Transgenerational Inheritance of Small RNAs in C. elegans. Cell 158: 277–287

Rechavi O & Lev I (2017) Principles of Transgenerational Small RNA Inheritance in Caenorhabditis elegans. Curr. Biol. 27: R720–R730

Rechavi O, Minevich G & Hobert O (2011) Transgenerational inheritance of an acquired small RNA-based antiviral response in C. elegans. Cell 147: 1248–56

Seth M, Shirayama M, Gu W, Ishidate T, Conte D & Mello CC (2013) The C. elegans CSR-1 Argonaute Pathway Counteracts Epigenetic Silencing to Promote Germline Gene Expression. Dev. Cell 27: 656–663

Seth M, Shirayama M, Tang W, Shen E-Z, Tu S, Lee H-C, Weng Z & Mello CC (2018) The Coding Regions of Germline mRNAs Confer Sensitivity to Argonaute Regulation in C. elegans. Cell Rep. 22: 2254–2264

Shen E-Z, Chen H, Ozturk AR, Tu S, Shirayama M, Tang W, Ding Y-H, Dai S-Y, Weng Z & Mello CC (2018) Identification of piRNA Binding Sites Reveals the Argonaute Regulatory Landscape of the C. elegans Germline. Cell 172: 937–951.e18

Shirayama M, Seth M, Lee HC, Gu W, Ishidate T, Conte D, Mello CC, Masaki Shirayama, 1, 2 Meetu Seth, 1 Heng-Chi Lee, 1 Weifeng Gu, 1 Takao Ishidate, 1, 2 Darryl Conte, Jr. 1 & and Craig C. Mello 1, 2 1 Program (2012) piRNAs Initiate an Epigenetic Memory of Nonself RNA in the C. elegans Germline. Cell 150: 65–77

Spracklin G, Fields B, Wan G, Vijayendran D, Wallig A, Shukla A & Kennedy S (2017) Identification and Characterization of C. elegans RNAi Inheritance Machinery. Genetics: 1–19

Towbin BDD, González-Aguilera C, Sack R, Gaidatzis D, Kalck V, Meister P, Askjaer P & Gasser SMM (2012) Step-wise methylation of histone H3K9 positions heterochromatin at the nuclear periphery. Cell 150: 934–947

Tyc KM, Nabih A, Wu MZ, Wedeles CJ, Sobotka JA & Claycomb JM (2017) The Conserved Intron Binding Protein EMB-4 Plays Differential Roles in Germline Small RNA Pathways of C. elegans. Dev. Cell 42: 256–270.e6

Vasale JJ, Gu W, Thivierge C, Batista PJ, Claycomb JM, Youngman EM, Duchaine TF, Mello CC & Conte D (2010) Sequential rounds of RNA-dependent RNA transcription drive endogenous small-RNA biogenesis in the ERGO-1/Argonaute pathway. Proc. Natl. Acad. Sci. U. S. A. 107: 3582–7

Verdel A, Jia S, Gerber S, Sugiyama T, Gygi S, Grewal SIS & Moazed D (2004) RNAi-Mediated Targeting of Heterochromatin by the RITS Complex. Science (80-.). 303: 672–676

Weiser NE, Yang DX, Feng S, Kalinava N, Brown KC, Khanikar J, Freeberg MA, Snyder MJ, Csankovszki G, Chan RC, Gu SG, Montgomery TA, Jacobsen SE & Kim JK (2017) MORC-1 Integrates Nuclear RNAi and Transgenerational Chromatin Architecture to Promote Germline Immortality. Dev. Cell 41: 408–423.e7

Zhang C, Montgomery TA, Gabel HW, Fischer SEJ, Phillips CM, Fahlgren N, Sullivan CM, Carrington JC & Ruvkun G (2011) mut-16 and other mutator class genes modulate 22G and 26G siRNA pathways in Caenorhabditis elegans. Proc. Natl. Acad. Sci. U. S. A. 108: 1201–8

Zhang D, Tu S, Stubna M, Wu W-S, Huang W-C, Weng Z & Lee H-C (2018) The piRNA targeting rules and the resistance to piRNA silencing in endogenous genes. Science 359: 587–592

